# Motor cortex can modulate somatosensory processing via cortico-thalamo-cortical pathway

**DOI:** 10.1101/366567

**Authors:** Michael Lohse, Matthew Cooper, Elie Sader, Antonia Langfelder, Martin Kahn, Luke Baxter, Julian Bartram, James W. Phillips, A. Louise Upton, Edward O. Mann

**Author notes:** These authors contributed equally to the work (A. Louise Upton, Edward O. Mann).

## Abstract

The somatosensory and motor systems are intricately linked, providing several routes for the sensorimotor interactions necessary for haptic processing. Here, we used electrical and optogenetic stimulation to study the circuits that enable primary motor cortex (M1) to exert top-down modulation of whisker-evoked responses, at the levels of brain stem, thalamus and somatosensory cortex (S1). We find that activation of M1 drives somatosensory responsive cells at all levels, and that this excitation is followed by a period of tactile suppression, which gradually increases in strength along the ascending somatosensory pathway. Using optogenetic stimulation in the layer-specific Cre driver lines, we find that activation of layer VI cortico-thalamic neurons is sufficient to drive spiking in higher order thalamus, and that this is reliably followed by excitation of S1, suggesting a cross-modal cortico-thalamo-cortical pathway. Cortico-thalamic excitation predicts the degree of subsequent tactile suppression, consistent with a strong role for thalamic circuits in the expression of inhibitory sensorimotor interactions. These results provide evidence of a role for M1 in dynamic modulation of S1, largely under cortico-thalamic control.

## Introduction

The integration of ascending somatosensory input with descending motor output is thought to be important for differentiating self- and externally-initiated tactile stimulation^1^, and interpreting somatosensory input during active touch. One would expect some facilitation of somatosensory responses during goal-directed movements^2^, but movement is generally associated with a suppression of tactile responses, including diminished responses to whisker stimulation in actively whisking rats^3^. The cortico-cortical circuits mediating sensorimotor interactions have been studied intensively in the rodent vibrissal system^4-7^, and it is has generally been found that exogenous activation of motor cortex acts to enhance somatosensory responses^8,9^. In contrast, it has been suggested that suppression of somatosensory responses can be established at the level of the brain stem, via inhibitory projections from the interpolaris subdivision of spinal trigeminal nucleus (SpVi)^3,10^. Less focus has been placed on physiological and behavioural role of motor cortico-thalamic contributions to somatosensory processing. The motor cortex has known excitatory projections to the somatosensory thalamus and ventral thalamus, and dorsal thalamus is modulated heavily by inhibitory nuclei in the ventral thalamus, principally zona incerta (ZI) and the thalamic reticular nucleus (TRN), providing the opportunity for dynamic gating of tactile processing. One type of engagement of motor cortex with the inhibitory circuitry of ventral thalamus was demonstrated by Urbain and Deschênes^11^, who showed that motor cortex interacts with the inhibitory circuitry within ZI, leading to fast inhibition of ZI outputs following motor cortical activation^11^. This suggests a physiological role for motor cortex projections in inhibitory and disinhibitory sensorimotor interactions involving dorsal thalamus.

The anatomical cortico-thalamic connectivity of rodent motor cortex is well understood. Motor cortex layer V projects directly to the higher order somatosensory thalamus (posterior medial nucleus; PoM), and more sparsely to cells in the somatosensory (vibrissal) first order nucleus (ventro-postero-medial nucleus; VPM)^11,12^. Furthermore, it was also shown recently that layer VI cortico-thalamic cells in motor cortex excite cells in PoM, and possibly project to VPM, as well as cortico-cortical layer VI cells^13^. In the canonical cortico-thalamic circuitry (derived from studying sensory cortico-thalamic projections), layer V is assumed to provide driving inputs to mostly higher order (e.g. PoM) thalamus, while layer VI provides modulatory inputs to the first order (e.g. VPM) thalamus from which it receives thalamo-cortical information. The nature of the differential physiological contribution of layer V and VI of motor cortex onto thalamus is poorly understood, and the physiological role of motor cortex in somatosensory processing in VPM and PoM *in vivo* during somatosensory stimulation remains unknown. Furthermore, the connectivity between motor cortex and somatosensory thalamus suggests a possible trans-thalamic pathway between motor and somatosensory cortices. Such trans-thalamic pathways have previously been found between primary and secondary sensory cortices^14,15^; however, it is currently unknown if such cortico-thalamo-cortical pathways exist between cortices processing different modalities, such as a motor and sensory system.

Here we study sensorimotor interactions in the mouse vibrissal system, and show that activation of motor cortex can drive both excitatory and inhibitory cells in cortex, as well as the thalamus and brain stem. This motor cortical drive is followed by a suppression of somatosensory responses at all three levels of the somatosensory system, but with progressive increase in the strength of suppression of somatosensory responses along the ascending somatosensory pathway, and almost full suppression apparent at the level of thalamus. Though layer V drives VPM and PoM, we also find that cortico-thalamic layer VI cells of motor cortex strongly drive PoM and to a lesser degree VPM. Stimulation of layer VI cortico-thalamic neurons drives responses in both thalamus and cortex, in a temporal sequence, with thalamus being activated before cortex, suggesting a cortico-thalamo-cortical pathway. In the thalamus, but not in the cortex, suppression of somatosensory responses is dependent on the strength of responses to stimulation of M1 layer VI (and to a lesser extent layer V). This suggests a critical role for cortico-thalamic drive in suppression of thalamic and cortical somatosensory responses following motor cortical output.

## Results

### Motor cortex can excite primary somatosensory cortex and subsequently inhibit somatosensory responses

The role of motor cortical activation in modulating spontaneous and whisker-evoked activity was initially explored in S1 of anaesthetised adult mice. Whole-cell patch-clamp recordings were obtained from S1 layer II/III neurons, and we recorded the responses to a combination of multi-whisker deflection and/or intracortical microstimulation of infragranular M1.

As expected, whisker stimulation evoked reliable EPSPs (Figure 1A&B). Electrical stimulation of M1 led to activation of S1 followed by a suppression of spontaneous activity for 100 – 150 ms (see Figure 1C). This decrease in spontaneous activity resembled a DOWN state of the cortical slow oscillation, which has previously been shown to enhance sensory responses^16^. Surprisingly, when M1 stimulation preceded whisker stimulation, the sensory-evoked EPSP was depressed (whisker-driven EPSP post motor cortex activation: 50 ms: *t(*4) = −5.49, *p* = 0.005, 75 ms: +*t(*5) = −2.50, *p* = 0.0543, 100-250 ms: *p* > 0.05) (Figure 1A-D). To confirm that the suppression induced by M1 stimulation could not be attributed to a switch to a cortical DOWN state under our anaesthetic protocol, we compared the whisker-evoked EPSPs that occurred during spontaneous UP and DOWN states. Whisker responses were enhanced in DOWN vs UP states (*p* < 0.001, *n* = 36; Wilcoxon test), and not suppressed as seen following M1 stimulation, which appears to dissociate M1-induced inhibition from processes occurring during a natural DOWN state (Figure 1E-H).

**Figure 1:**
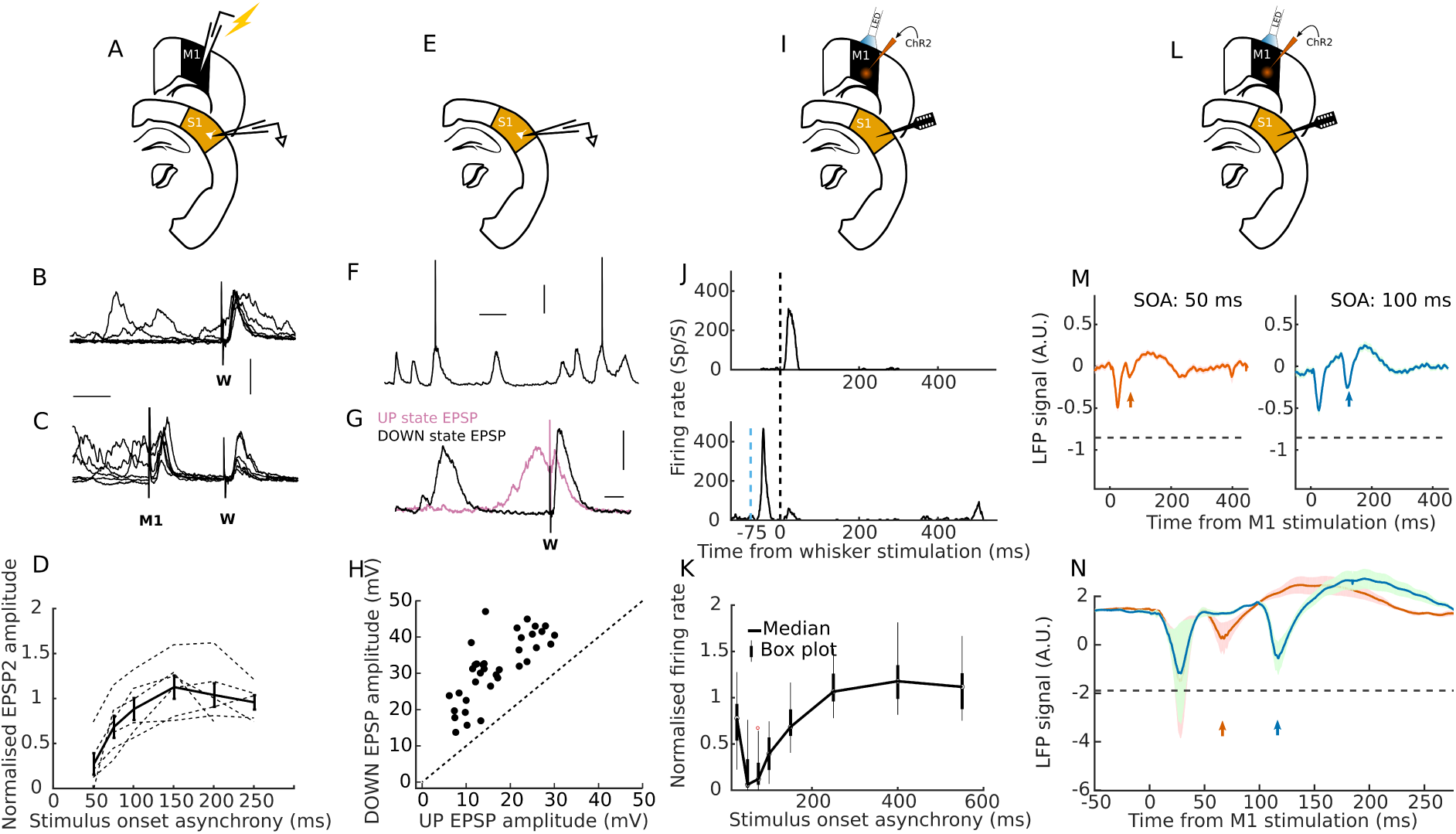
Motor cortical-induced excitation and subsequent inhibition in primary somatosensory cortex *in vivo*. A-D) Whole-cell recording from an S1 LII/III cell under ketamine/medetomidine anaesthesia, showing A) schematic illustrating recording and stimulation setup for B-D, B) example of whisker-evoked EPSPs, and C) example of responses to electrical activation of M1, which is associated with a suppression of subsequent whisker-evoked EPSPs. Calibration bars: 50 ms and 10 mV. D) Summary of M1-induced suppression of subsequent whisker-evoked EPSPs in S1 LII/III cells (normalised to EPSP amplitude to whisker stimulation without preceding M1 activation). Solid line: mean ± s.e.m. E-H) *In vivo* subthreshold dynamics of S1 LII/III cells displaying spontaneous UP and DOWN states. E) Schematic illustrating recording setup for F-H. F) Example of naturally occurring UP and DOWN states. Calibration bars: 200 ms and 20 mV. G) Example of whisker-evoked EPSPs during spontaneous UP (purple) or DOWN (black) states. Calibration bars: 10 ms and 2 mV. H) Whisker responses (EPSP amplitude) are larger during DOWN states relative to UP states. I-K) Optogenetic activation of CaMKII-expressing neurons in M1 induces responses (MUA firing rate) in S1 followed by prolonged inhibition of whisker-evoked MUA responses. I) Schematic illustrating recording and stimulation setup for J&K. J) Example whisker-evoked response in S1 MUA (top) and M1-induced response (bottom) in S1 MUA recordings. K) Summary of M1-induced suppression of subsequent whisker-evoked MUA responses in S1 (normalised to MUA response to whisker stimulation without preceding M1 activation). L-N) Optogenetic activation of M1 induces responses in LFPs in S1 followed by prolonged inhibition of whisker induced LFP responses. M) Schematic illustrating recording setup for M&N. M) Example whisker induced response in S1 LFPs following M1 activation. N) Mean LFP traces (across animals) of M1-induced suppression of subsequent incoming whisker induced LFP responses in S1 (dashed horizontal line represents LFP amplitude to non-modulated (no M1 stimulation) whisker stimulation). Shaded area represents ± s.e.m.

A similar temporal profile of suppression of somatosensory responses could also be found when using paired whisker deflections (Supplementary Figure 1I-J). Both intramodal and cross-modal interactions were also observed in suprathreshold MUA responses, recorded with multielectrode linear probes (data not shown). Together, this suggests that electrical stimulation of M1 can suppress both sub- and suprathreshold somatosensory responses, and that the circuitry mediating these delayed inhibitory effects may also be recruited by sensory input.

Electrical simulation in M1 antidromically activates afferent projections, including those coming from S1 and thalamus. We therefore wanted to determine whether optogenetic activation of M1 could also suppress somatosensory responses. For optogenetic transduction, AAV-CaMKII-ChR2-YFP was injected unilaterally in to the infragranular layers of M1. To confirm functional expression, we obtained whole-cell recordings in *ex vivo* brain slices of S1, and found that optogenetic activation of M1 axons in S1 could drive responses in both inhibitory and excitatory neurons, as previously reported (Supplementary Figure 1). We did not find any evidence for an interaction between M1-evoked responses and EPSPs evoked by local electrical stimulation (Supplementary Figure 1), which may be due to the severing of local and/or long-range projections in this reduced *ex vivo* preparation. However, these experiments confirmed that we could use optogenetics successfully to drive M1 outputs, and we proceeded to use optogenetic stimulation for all further experiments.

To assess whether optogenetic stimulation of M1 could evoke an excitatory-inhibitory sequence in CaMKII-ChR2-injected mice *in vivo*, we recorded extracellular MUA and LFP responses from S1 across all layers using multielectrode linear probes. In S1 *in vivo*, 58/64 (90.6 %) recorded MUAs responsive to whisker stimulation *(p* < 0.005 as inclusion criterion; t test) could also be significantly driven *(p* < 0.05; t test) by optogenetic stimulation of infragranular CaMKII cells expressing ChR2-YFP in M1. The evoked activity was followed by a suppressive period in both the MUA and LFP, as was previously seen in the subthreshold dynamics (EPSPs) of S1 cells (MUA: 25 ms - 150 ms post motor cortex activation *p* < 0.001, *n* = 64 MUAs (4 mice); Wilcoxon test; Figure 1I-N). The extracellular MUA recordings furthermore revealed an extended period of slight hyperexcitability (200-550 ms post motor cortex activation*, p* < 0.001, *n* = 64; Wilcoxon test).

These results demonstrate a triphasic excitation-inhibition-excitation pattern in S1 induced by motor cortex stimulation, with a period of tactile suppression that appears unrelated to the concurrent modulation of cortical UP and DOWN states.

### Motor cortex induced inhibition of somatosensory responses is weakly present in the somatosensory brain stem and robustly implemented from thalamus onwards

While M1 sends direct cortico-cortical projections to S1, M1 corticofugal projections may contribute to activation of S1 via relay pathways, and establish the subsequent suppression of activity earlier in the somatosensory pathway. Therefore, having recorded the S1 responses in the CaMKII-ChR2 injected mice, we sequentially recorded from brain stem (trigeminal nuclei) and thalamus (recordings predominantly targeted to PoM), in order to explore the stage at which these activity patterns emerged (Figure 2).

**Figure 2:**
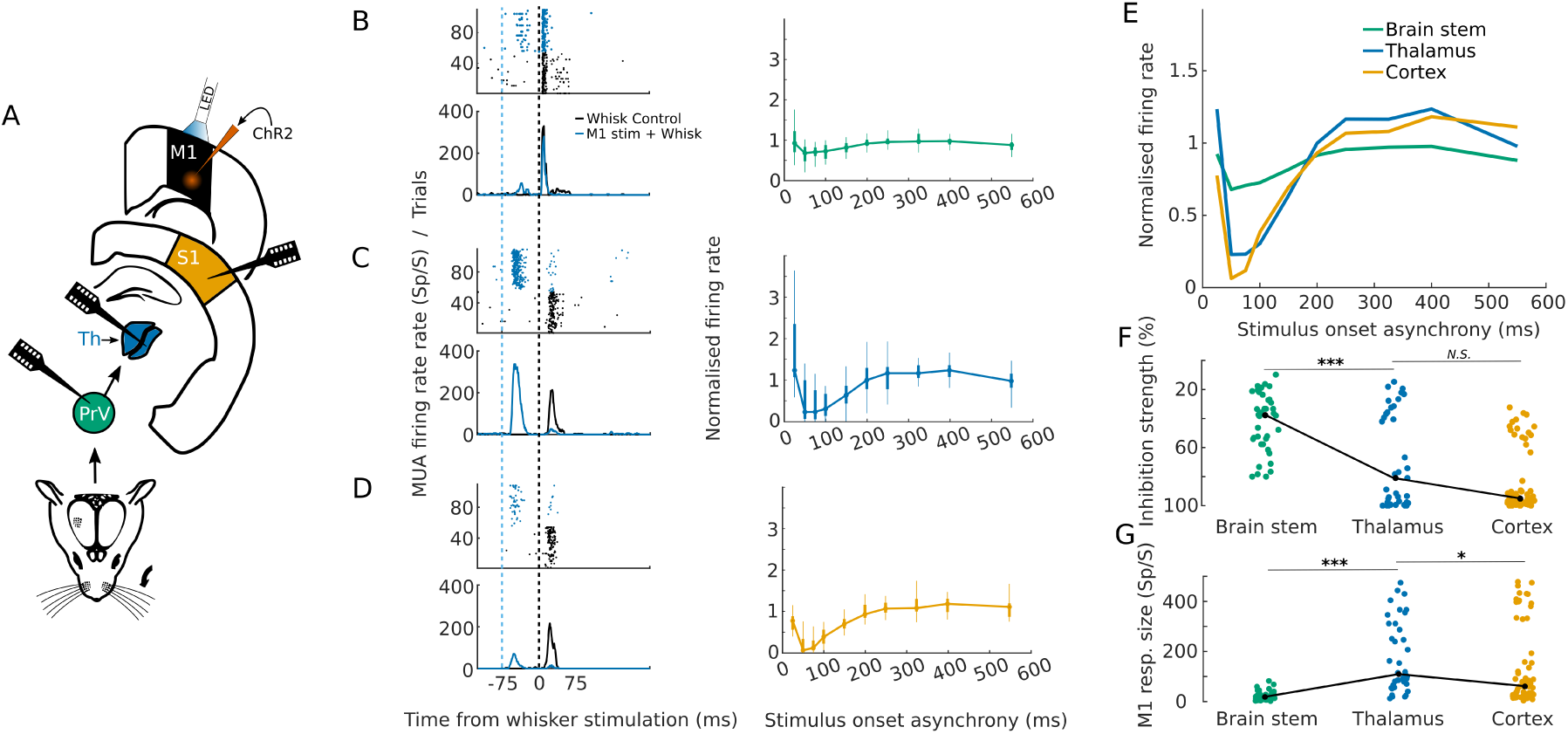
Increasing M1-induced inhibition along the somatosensory pathway. A) Schematic illustrating recording and stimulation setup. B-D) Modulation by ChR2 activation in M1 CaMKII-expressing cells at different levels of the somatosensory pathway. Left: Example raster plots showing responses to whisker stimulation with and without prior M1 activation (upper plots), and the corresponding spike time histograms (lower plots). Right: Summary of M1-induced suppression, with activity averaged (median) over all MUAs recorded within each region in order to assess the overall M1-induced inhibition at each level in the somatosensory pathway (4 mice): B) Brain stem (trigeminal nuclei, *n* = 38), C) Thalamus (VPM/PoM, *n* = 39), D) Cortex (S1, *n* = 64). E) Summary showing the overlay of median modulation from B-D. F) Increasing maximum inhibition, showing responses for individual MUA with median overlaid (solid line). G) Strength of M1 stimulation induced firing rate in whisker-responsive MUAs recorded in the somatosensory brain stem, thalamus and cortex, with median overlaid (solid line) *(*p* < 0.05, *\*\*\*p* < 0.001, *N.S.:* p > 0.05, Mann-Whitney U test)

Firstly, we found that MUAs responsive to whisker stimulation in somatosensory thalamus could also be reliably driven by activation of CaMKII-expressing cells in M1. We found that 24/35 (68.6 %) recorded MUAs responsive to whisker stimulation in somatosensory thalamus were significantly *(p* < 0.05; t test) driven by M1 stimulation. In the somatosensory brainstem, a smaller fraction of MUAs responsive to whisker stimulation were found to be significantly driven by M1 stimulation (18/34; 52.9%). When assessing the strength of the drive we found that MUAs in S1 and somatosensory thalamus were driven significantly stronger than the somatosensory brainstem by M1 stimulation *(p* < 0.001; Mann-Whiney U test; Figure 2A-D&G), and that MUAs in thalamus were driven slightly, but significantly, more strongly than those in S1 *(p* = 0.041; Mann-Whiney U test; Figure 2A-D&G).

Secondly, we found that the whisker suppressive effect following motor cortex activation (measured as % inhibition relative to control response at the stimulus onset asynchrony (SOA) with maximal inhibition for each mouse) was not only present in S1 (95.2 % of median inhibition*; p <* 0.001; Wilcoxon test), was also present at the level of the somatosensory brain stem (37.7 % median inhibition; *p <* 0.001; Wilcoxon test), and robustly present at the level of the thalamus (81 % median inhibition; *p <* 0.001; Wilcoxon test; Figure 2B-F). The suppressive effect was significantly less in brain stem compared to thalamus and S1 *(p <* 0.001 and *p <* 0.001, respectively; Mann-Whiney U test), while the effects observed in thalamus and S1 were not significantly different *(p =* 0.059; Mann-Whiney U test). This shows that stimulation of M1 leads to a robust activation and subsequent whisker response suppression in somatosensory thalamus, and suggests a primarily thalamic origin of the delayed suppression of responses seen in S1.

### Motor cortico-thalamic projections from layer V and Layer VI can both drive spiking activity in somatosensory thalamus

Cortex has two major corticofugal projections from layer V and layer VI. In accordance with canonical understanding of cortico-thalamic circuitry from sensory systems, layer V is thought to provide driving input forward to the next higher order thalamic nucleus (e.g. S1 to PoM), while layer VI is supposed to only provide modulatory information back to the primary thalamic relay nucleus (e.g. S1 to VPM)^17^. A subset of layer V cells (mostly located in layer Va) and most (if not all) layer VI cortico-thalamic cells can be targeted selectively using the transgenic RBP4-cre and NTSR1-cre mouse lines, respectively^18,19^. In order to understand whether these two major corticofugal outputs play a differential role in the M1-induced modulation of the somatosensory thalamo-cortical system, we injected floxed ChR2-YFP virus into M1 of these transgenic Cre driver lines, thus enabling layer-specific optogenetic activation.

To first explore the direct functional connectivity between M1 and somatosensory thalamus, we combined *ex vivo* whole-cell patch-clamp recordings in PoM with optical stimulation of ChR2-YFP expressing M1 layer V or layer VI cortico-thalamic axons, in RBP4-Cre or NTSR1-Cre mice, respectively (Figure 3A). We confined our whole-cell recordings to PoM, as we could identify very few ChR2-YFP expressing axons in VPM in our *ex vivo* slices. We found synaptic responses following activation of both types of axon (Figure 3B&C), with the responses to light activation of layer V cortico-thalamic axons having a shorter 20-80 % rise time (median rise times [IQR] following stimulation of layer V vs VI cortico-thalamic axons: 1.3 [1.0, 1.7] vs 2.1 [1.6, 4.6] ms; *p* = 0.01, n = 8 and 25, respectively; Mann-Whitney U test; Figure 3D), and a steeper initial EPSP slope (median EPSP slopes: 3.0 [1.4, 1.8] vs 0.86 [0.35, 1.9] V/s; *p* = 0.017; Mann-Whitney U test). Both of these features could be consistent with layer V projections forming large proximal synapses on thalamic relay cells. However, there appeared to be a higher probability of finding an evoked response following light activation of layer VI cortico-thalamic axons (proportion of detected responses following layer V vs layer VI cortico-thalamic axons: 8/16 vs 25/30), and this evoked response was at least as likely to drive postsynaptic spiking (Figure 3E).

**Figure 3:**
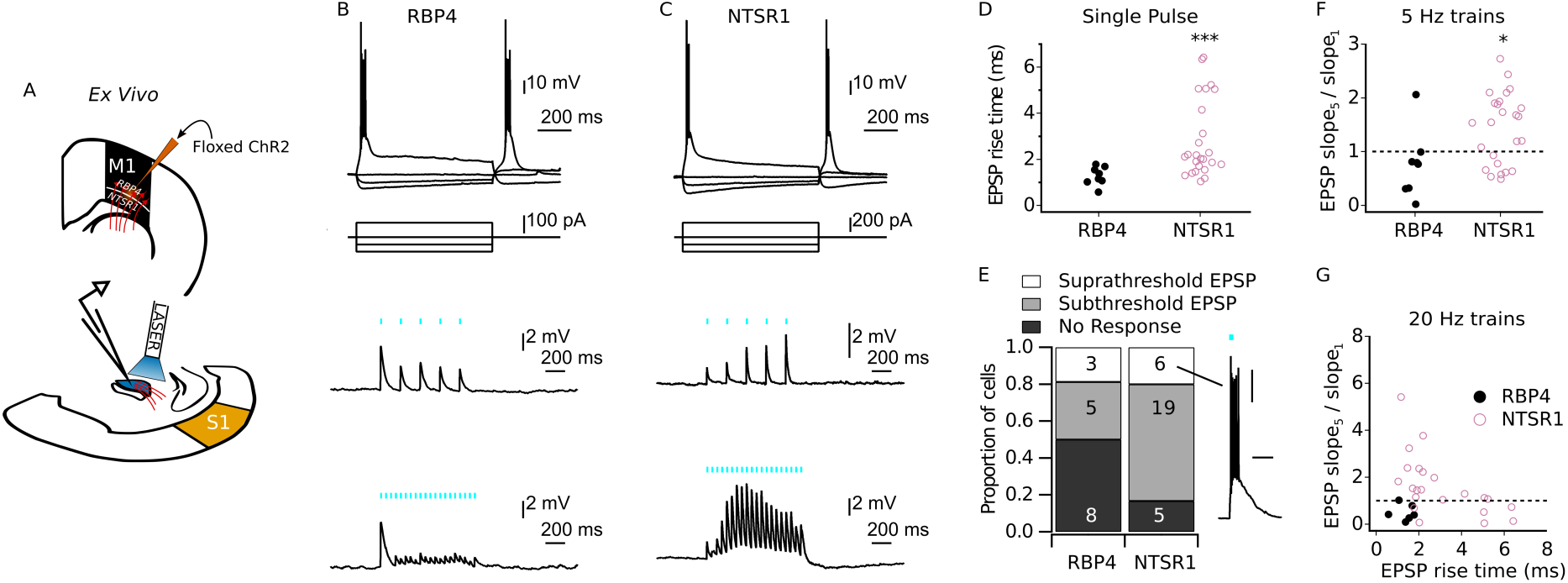
M1 layer V and VI differentially excite cells in PoM *ex vivo.* A) Schematic illustrating *ex vivo* recording and stimulation setup. B-C) Whole-cell current-clamp recordings were made from PoM neurons in slices from RBP4-Cre (B) and NTSR1-Cre mice (C), which had received M1 injections of floxed ChR2-YFP. All neurons displayed rebound burst firing, typical of thalamic relay neurons (top). Cortico-thalamic axons were stimulated with single 20 ms light pulses at 100 % laser power, or as light pulse trains at 5 Hz (middle) and 20 Hz (bottom) for 1 s. D) 20-80% rise times for EPSPs evoked by single light pulses. E) Proportion of cells recorded that displayed no response, subthreshold responses, and suprathreshold responses. F) Short-term plasticity of cortico-thalamic transmission with 5 Hz trains, measured as the ratio of the EPSP slope following the 1^st^ and 5^th^ light pulse. G) EPSP slope ratio for 20 Hz trains plotted against EPSP rise time, showing the diversity of response profiles recorded in response to stimulation of layer VI cortico-thalamic axons. (*** *p* = 0.001, * *p* < 0.05, Mann-Whitney U test)

Cortico-thalamic inputs from layer V and layer VI are also thought to be distinguished by their short-term plasticity, with layer V inputs showing short-term depression and layer VI inputs showing facilitation. In most cells, we were also able to examine the responses to 1 s light pulse trains at 5 and 20 Hz. For the 20 Hz trains, the responses often waxed and waned across 1 s train, and we therefore analysed the ratio of the EPSP slope for 5^th^ versus 1^st^ pulse, in each case. This EPSP slope ratio was significantly higher with activation of layer VI versus layer V cortico-thalamic axons, during both the 5 Hz (median EPSP ratios [IQR] following stimulation of layer V vs layer VI cortico-thalamic axons: 0.78 [0.31, 0.95] vs 2.1 1.54 [0.72, 1.9]; *p* = 0.014, n = 8 and 25, respectively; Mann-Whitney U test; Figure 3F) and 20 Hz trains (0.40 [0.22, 0.84] vs 1.4 [0.72, 2.3]; *p* = 0.029, n = 6 and 22, respectively; Mann-Whitney U test; see Figure 3G), but there were many responses to activation of layer VI cortico-thalamic axons that displayed either little short-term plasticity or depression (Figure 3F & G). Overall, while the number of responses recorded following activation of layer V cortico-thalamic axons was relatively small, they appeared to display the relatively homogenous properties expected of a driver input. In contrast, the responses evoked by activation of layer VI cortico-thalamic axons were diverse, and there did not appear to be a clear cluster of responses with slow rise times and short-term facilitation, which one might expect from a typical modulator input.

To examine the differential contribution of M1 layer V and layer VI projections to excitatory responses in somatosensory thalamo-cortical circuits *in vivo*, we recorded MUA responses in VPM, PoM and S1 following optical stimulation of M1 layer V (RBP4) or layer VI (NTSR1) neurons, and compared these to the responses evoked by whisker stimulation (Figure 4). We identified our thalamic recording sites as PoM or VPM by a combination of histologically identifying tract traces (DiI staining), and the latency of the sensory response^20^. The histogram of whisker response latencies showed a clear bimodal distribution, with the trough at 20 ms, allowing us to separate putative VPM cells (< 20 ms latency) from PoM cells (> 20 ms latency) (Figure 4A-D). Using this classification, we found that stimulating layer V cortico-thalamic neurons in motor cortex evoked significant *(p* < 0.05) spiking activity in 21/87 (24 %) whisker-responsive MUAs in S1, 46/71 (64.8 %) in VPM, and 57/82 (69.5 %) in PoM (Figure 4). Similarly, activation of layer VI cortico-thalamic neurons was found to excite 24/57 (42%) MUAs in S1, 16/38 (42 %) in VPM, and 48/68 (70.5 %) in PoM. M1 layer V had stronger connectivity (induced firing rate) with VPM compared to layer VI *(p* < 0.005; Mann-Whiney U test), while the strengths of excitation (induced firing rate) from layer V and layer VI were similar for PoM responses *(p* = 0.21; Mann-Whiney U test). PoM was excited (induced firing rate) to a greater degree relative to VPM when stimulating either layer V *(p* < 0.001; Mann-Whiney U test) or layer VI *(p* < 0.001; Mann-Whiney U test; Figure 4).

**Figure 4:**
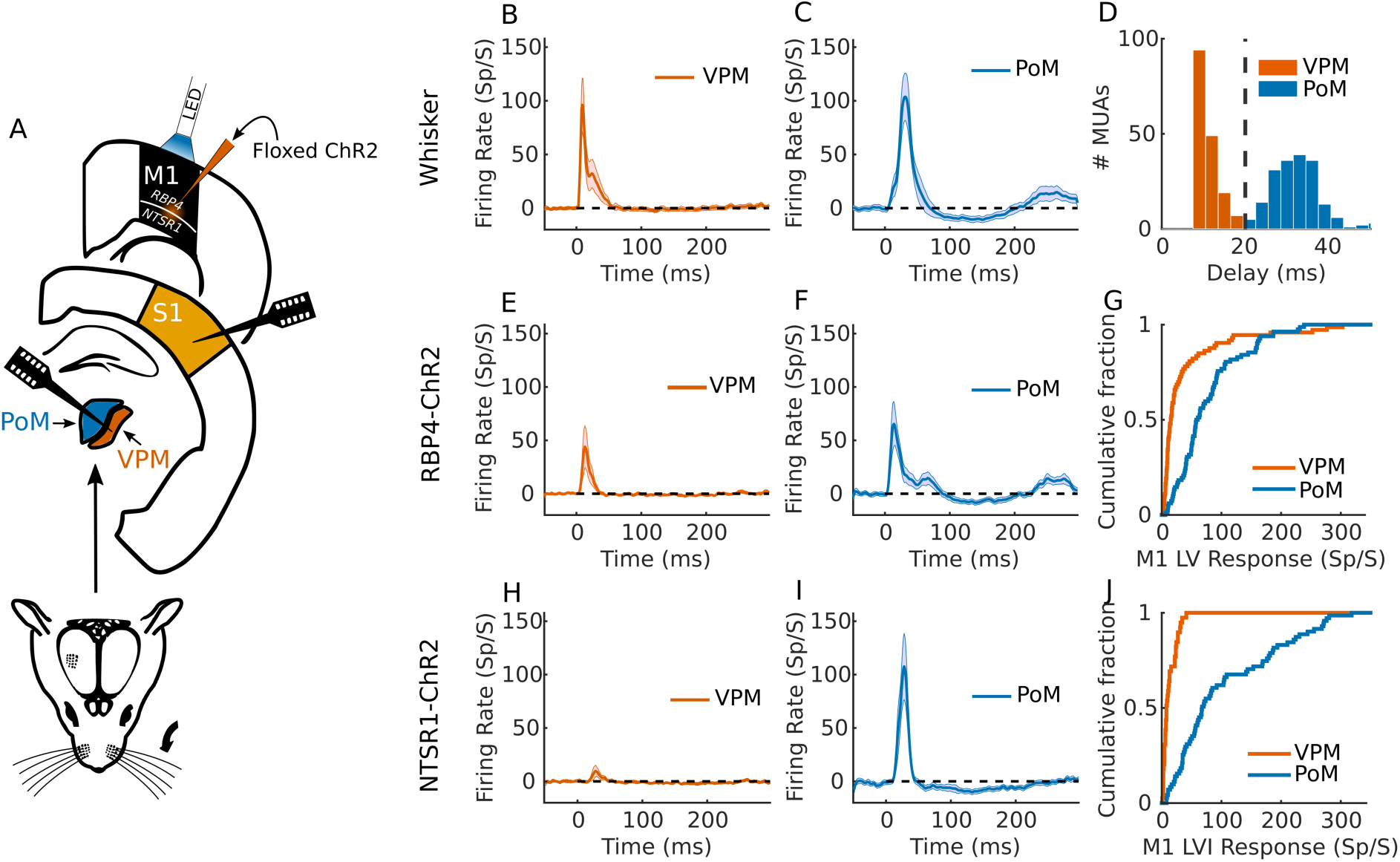
Layer-specific motor cortico-thalamic drive of the somatosensory thalamo-cortical system. A) Schematic illustrating recording and stimulation setup. B-C) Mean response (across MUAs) to whisker stimulation in VPM (B), and PoM (C). D) Response latencies (delay to peak firing rate) in somatosensory thalamus, used to parse out VPM and PoM recordings - PoM and VPM MUAs are separated at peak latency responses of 20 ms (dashed line). E-G) M1 layer V (RBP4) induced response characteristics in somatosensory thalamus and S1. E-F) Mean response (across MUA) to M1 layer V stimulation in VPM (E), and PoM (F). G) Cumulative histogram of response magnitude to M1 layer V (RBP4) optogenetic stimulation (MUA firing rate) in VPM and PoM. H-J) M1 layer VI (NTSR1) induced response characteristics in somatosensory thalamus H-I) Mean response (across MUAs) to M1 layer VI stimulation in VPM (H), and PoM (I). J) Cumulative histogram of response magnitude (MUA firing rate) to M1 layer VI (NTSR1) optogenetic stimulation (MUA firing rate) in VPM and PoM. In panels B-I, shaded area represents 95 % confidence intervals.

Given the divergent cortico-cortical and corticofugal projections from RBP4+ layer V neurons, the robust responses observed across the somatosensory thalamo-cortical circuit are not surprising. Indeed, consistent with its role as a thalamic driver, stimulation of M1 layer V produced shorter response times in thalamus relative to layer VI (M1-layer V → VPM vs M1-layer VI → VPM, *p* = 0.001; M1-layer V → PoM vs M1-layer VI → PoM, *p* < 0.001). Some of these difference in response delay could be due to light dispersion, leading to stronger and faster activation of layer V located more superficially in the neocortical layers cells, but they are likely to be due predominantly to faster axon conduction velocities^21^ and faster EPSP rise times (as confirmed *ex vivo* in Figure 3). However, despite this fast cortico-thalamic drive, current source density (CSD) analysis showed that S1 activation by M1 layer V was relatively weak and dispersed (Figure 5), suggesting that that M1 layer V cortico-cortical and corticofugal projections do not cooperate to drive S1 activity.

**Figure 5:**
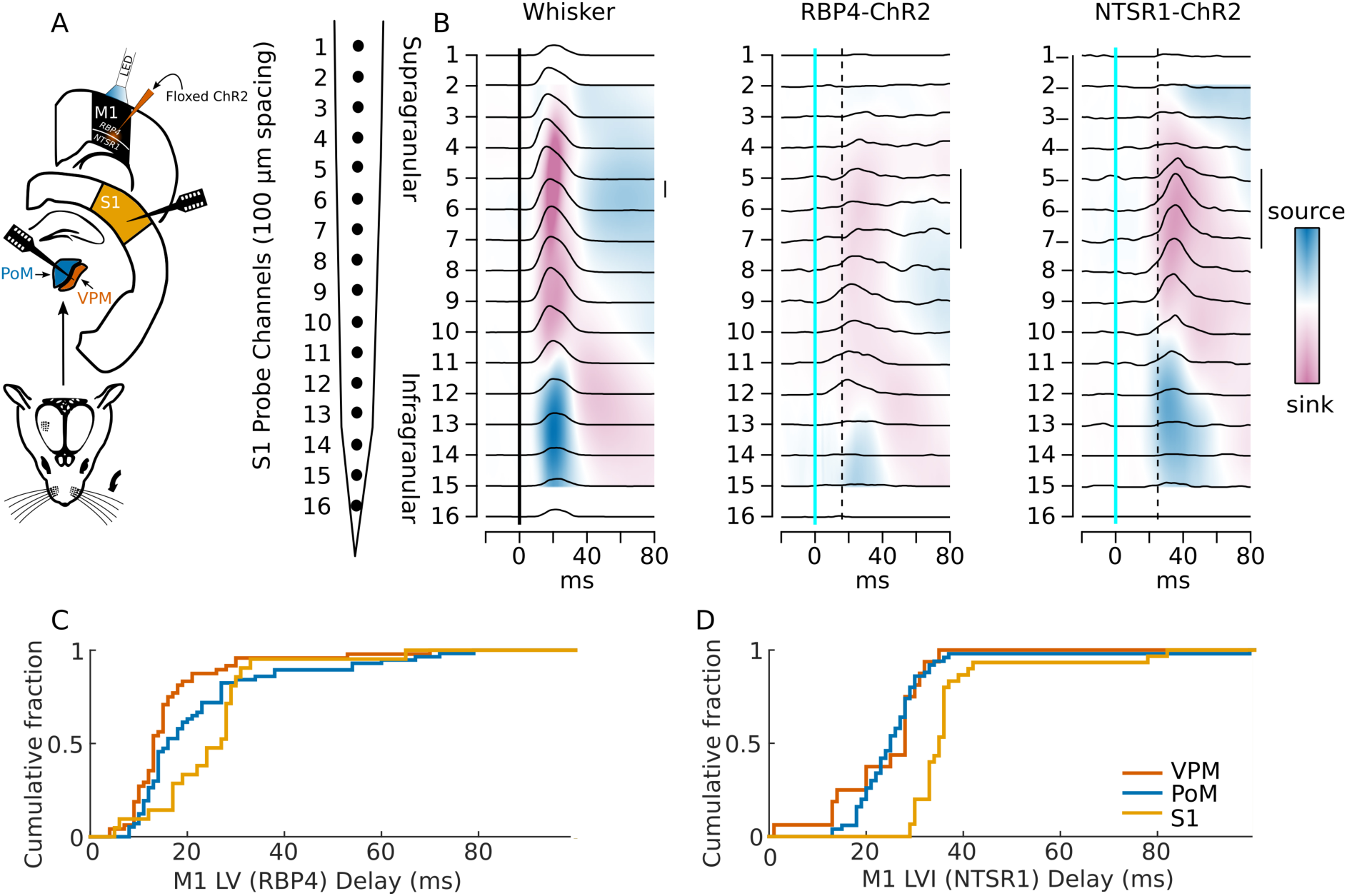
Temporal progression of excitation of the somatosensory thalamo-cortical system following M1 stimulation provides evidence for a cortico-thalamo-cortical pathway from M1 to S1. A) Schematic illustrating recording and stimulation setup. B) Current source density (CSD) analyses of S1 responses to whisker stimulation (left), M1 layer V (RBP4) stimulation (middle), or M1 layer VI (NTSR1) (right). S1 MUA responses for each channel are overlaid on top of CSD. Calibration bar =100 Sp/S. Solid vertical lines indicate stimulus onset and stippled vertical lines indicate median delay to peak response for PoM neurons following either M1 layer V or M1 layer VI stimulation. C-D) Cumulative histogram illustrating response latencies (delay to peak MUA firing rates) to M1 layer V (C) and M1 layer VI (D) stimulation across VPM, PoM and S1.

The activation of both somatosensory thalamus and S1 following activation of M1 layer VI cortico-thalamic neurons is perhaps more surprising, but consistent with the functional connectivity observed in *ex vivo* slices (Figure 3). As NTSR1-expressing layer VI cells in M1 only very weakly innervate layer V of M1 (Supplementary Figure 2; see also^13^), and have no long range cortico-cortical connections^18^, this appears to suggest a direct layer VI cortico-thalamic drive of neurons in somatosensory thalamus (Figure 4J). This result is supported by the finding that S1 activation by M1 layer VI stimulation is significantly delayed (VPM and PoM vs S1: 7-10 ms delay) relative to somatosensory thalamus activation, Kruskal-Wallis test, *p* < 0.001, post-hoc Dunn-Sidak test: VPM vs PoM, *p* = 0.99, VPM vs S1, *p* < 0.001, PoM vs S1, *p* < 0.001 (Figure 5). Furthermore, CSD analysis of the M1 layer VI induced response across layers in S1 revealed a robust and locked response focused around the thalamo-cortical (high order) input layers, suggesting a reliable transfer of the M1 layer VI activation, potentially through the thalamus.

It is unclear whether this non-modulatory excitation of M1 layer VI onto somatosensory thalamic cells happens during normal functioning of the cortico-thalamic sensorimotor system, or whether it is a feature of the artificial activation of many axons synchronously releasing glutamate onto distal dendrites of a cell surrounded by ChR2 expressing layer VI motor cortical axons. However, the *ex vivo* results provide evidence for how the driven activity of somatosensory thalamic cells by M1 layer VI could arise monosynaptically when stimulating M1 layer VI optogenetically *in vivo*, and provides evidence for how M1 layer VI has the possibility of providing an excitatory spread of information through a cortico-thalamo-cortical pathway, from M1 layer VI to S1.

### Suppression of incoming somatosensory information depends on strength of M1 excitation in individual somatosensory thalamic units

We finally assessed the role of thalamus in establishing the delayed suppression of incoming somatosensory information following M1 layer V or layer VI stimulation. We find that both layer V and layer VI stimulation can inhibit subsequent somatosensory responses in both VPM (layer V: *p* < 0. 001, *n* = 73; layer VI: *p* < 0.001, *n* = 39; Wilcoxon test) and PoM (layer V: *p* < 0.001, *n* = 82; layer VI: *p* < 0.001, *n* = 71; Wilcoxon test), as well as S1 (layer V: *p* < 0.001, *n* = 88; layer VI: *p* < 0.001, *n* = 60; Wilcoxon test) (Figure 6). For both layer V and layer VI, somatosensory responses in PoM were significantly more suppressed compared to VPM (layer V: *p* < 0.001; layer VI: *p* = 0.002; Mann-Whitney U test). Layer V provided stronger inhibition of PoM than layer VI *(p* < 0.01; Mann-Whitney U test), while this difference could not be found in VPM *(p* = 0.43; Mann-Whitney U test).

**Figure 6:**
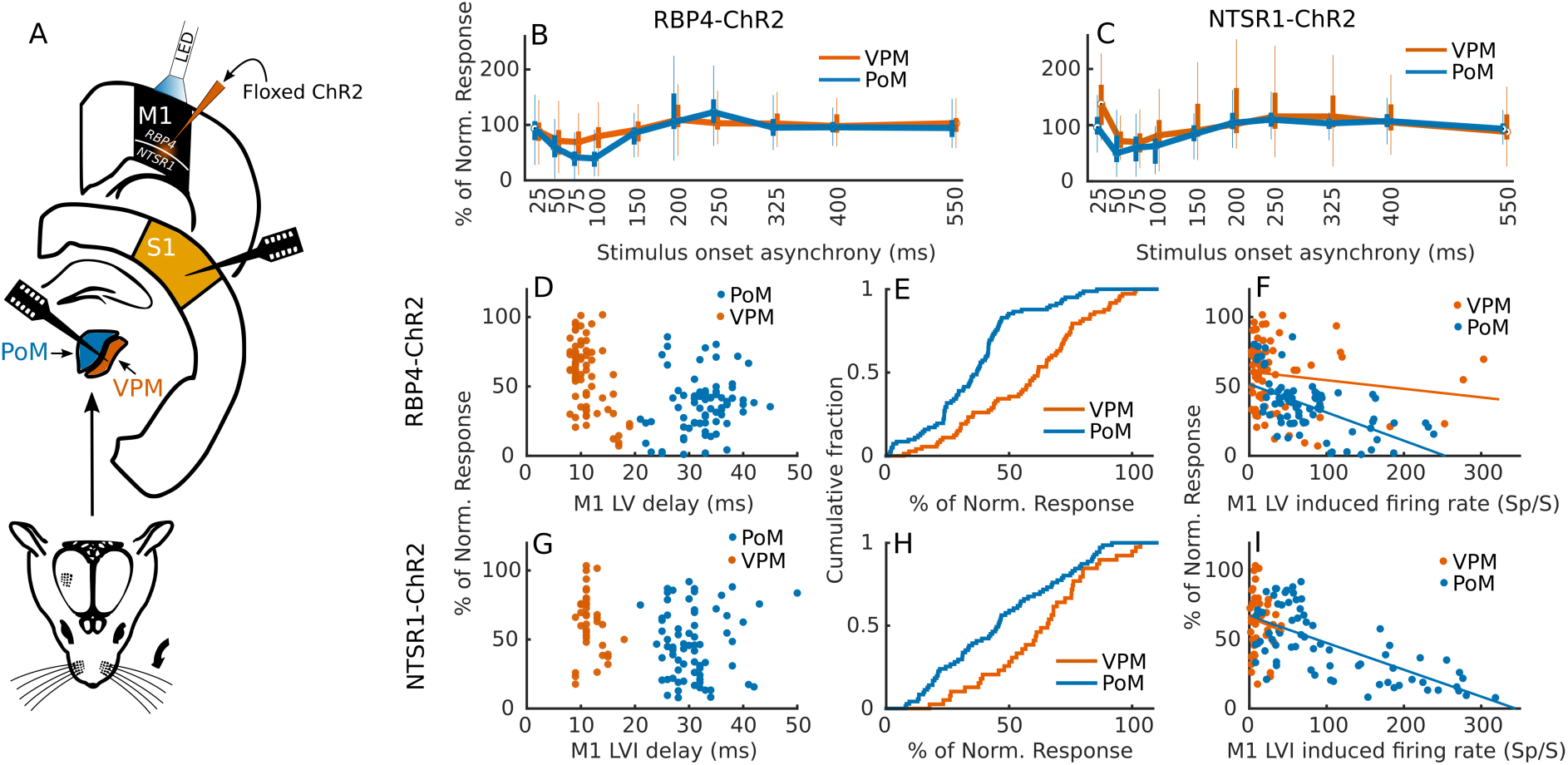
Motor cortex induced inhibition of thalamic somatosensory responses correlates with the degree of local thalamic excitation. A) Schematic illustrating recording and stimulation setup. B-C) Suppression (% firing rate relative to unmodulated whisker response) of incoming response to whisker deflection as a function of stimulus onset asynchrony between M1 layer V (B), or layer VI (C) stimulation and whisker stimulation. D-I) Analysis of inhibition observed across individual electrode sites. D&G) Maximum inhibition in VPM and PoM following M1 stimulation (percentage MUA firing rate relative to non-modulated whisker response) by layer V (RBP4; D) or layer VI (NTSR1; G) stimulation, plotted against the delay of whisker-evoked responses at the same site. E&H) Cumulative histogram illustrating maximum inhibition strength in VPM and PoM following stimulation of M1 layer V (E) or layer VI. (H). F&I) Relationship between excitation (M1-induced firing rate) and suppression of subsequent whisker response (maximum inhibition) in VPM and PoM after stimulation of M1 layer V (F) or layer VI (I).

Importantly, we found a strong positive relationship between excitation of a PoM multi-unitz and the amount of subsequent suppression of incoming somatosensory responses (Figure 6F & I). This relationship was present for stimulation of both layer V (*r* = −0.57*, p* < 0.001; Pearson’s Correlation) and layer VI (*r* = −0.68*, p* < 0.001; Pearson’s Correlation). This excitation-inhibition was inverted in S1, where multi-units responding strongly to M1 stimulation are almost unaffected by M1 stimulation, while multi-units responding weakly to M1 stimulation are more affected by M1 stimulation (layer V: *r* = 0.46*, p* < 0.001; layer VI: *r* = 0.28 *,p* = 0.03; Pearson’s Correlation). Re-examining our previous recordings, we found no relationship between M1-induced (CaMKII-positive cells) excitation and subsequent inhibition of whisker responses was found in the brain stem (*r* = 16, *p* = 0.34; Pearson’s Correlation). These results demonstrate that cortico-thalamic excitation is needed for a thalamic multi-unit, but not a cortical multi-unit, to suppress incoming somatosensory information, suggesting a strong thalamic component in establishing suppression of incoming somatosensory responses.

## Discussion

We have demonstrated that activation of infragranular layers V and VI of primary motor cortex can drive activity in somatosensory pathways, and subsequently suppress whisker-evoked responses for a period of ~100 ms. Selective activation of M1 NTSR1+ layer VI neurons was sufficient to induce this excitatory-inhibitory cascade, suggesting that at least one pathway of modulation is via M1 projections to somatosensory thalamus. Indeed, activation of either M1 RPB4+ layer V neurons or M1 NTSR1+ layer VI neurons evoked robust responses in PoM, with a significant correlation between the degree of local excitation and subsequent inhibition. The patterns of excitation and inhibition in the somatosensory thalamus appeared to be reflected in S1 responses, suggesting that M1 can interact with S1 via a cortico-thalamo-cortical pathway.

Cortico-thalamo-cortical pathways have previously been found between primary and secondary sensory cortical areas^14,15^, but such pathways between modalities remain to be established. Previous optogenetic mapping studies in *ex vivo* slices have suggested that M1 cortico-thalamic projections primarily act to excite neurons in somatosensory thalamus^13^, which we have confirmed here in both *ex vivo* slices and *in vivo.* While we found that activation of either M1 RBP4+ layer V or NTSR1+ layer VI neurons could drive activity in somatosensory thalamus, optogenetic activation of M1 NTSR1+ layer VI neurons evoked a clear and delayed sequence of thalamic and cortical activation, suggesting a cortico-thalamo-cortical circuit that crosses from motor to a sensory system. Given the proposed roles of layer V and layer VI cortico-thalamic neurons as drivers and modulators, respectively^17^, the ability of layer VI cortico-thalamic neurons to drive thalamo-cortical activity may be somewhat surprising. This could relate to nature of the optogenetic stimulation paradigm, which evokes synchronous activity across a population of neurons/axons. Indeed, examining the dynamics of optogenentically-evoked synaptic responses in thalamic neurons in *ex vivo* slices revealed many of the features expected of drivers and modulators, with EPSPs evoked by stimulation of M1 RBP4+ axons showing fast rise times and synaptic depression (driver), and those evoked by stimulation of M1 NTSR1+ axons showing slow rise times and/or synaptic facilitation (modulator)^17,22,23^. However, the synaptic dynamics for M1 NTSR1+-evoked responses were diverse, without a clear cluster of responses with both slow rise times and synaptic facilitation, and stimulation of M1 NTSR1+ axons was at least as likely to evoke spiking activity as stimulation of M1 RBP4+ axons. It may be that layer V and layer VI cortico-thalamic outputs have different roles depending on cortical origin and/or projection target^24^. While the optogenetic experiments presented here suggest a functional excitatory cortico-thalamo-cortical pathway between M1 and S1, how this pathway might be recruited during natural patterns of activity remains to be resolved.

The activation of infragranular layers V and VI of M1 was followed by a period of tactile suppression, with an amplitude and time course similar to that observed following intracortical microstimulation of M1 in monkeys^25^. Such tactile suppression is also observed during movement^26-29^, and is thought to inhibit responses to self-generated tactile stimulation. Surprisingly, previous studies examining the effects of M1 optogenetic stimulation on whisker-evoked responses in mice have found predominantly facilitatory effects^8,9^. This may be partly explained by the nature of the optogenetic stimulation paradigm, and sustained optogenetic activation of M1 appears to modulate sensory responses via cortico-cortical projections, and effects on network state^9^. However, a recent study used a light pulse stimulation paradigm equivalent to ours, and found enhancement of whisker-evoked responses for up to 50 ms after M1 simulation, with no evidence of suppression^8^. In this case, the ChR2 injection sites were more rostral than used here, and thus may have activated a different functional domain of M1^6^, with a potentially different role in sensorimotor processing.

Tactile suppression is likely to be mediated at multiple levels within sensorimotor systems, and may be effected by corollary motor commands and proprioceptive reafference^3,8,10,26-30^. Here, we find that activation of M1 NTSR1+ layer VI cortico-thalamic neurons induces an excitation-suppression sequence across thalamocotical circuits, suggesting that cortico-thalamic projections from M1 to somatosensory thalamus are sufficient to induce some aspects of tactile suppression. In this case, trans-thalamic excitation via PoM may recruit additional intracortical inhibitory mechanisms^31^. However, the predominant response to PoM stimulation *in vivo* is an enhancement of whisker responses^32^, and it seems likely that the suppression of cortical responses is largely inherited from thalamus. One major source of inhibition to sensory thalamus is supplied by ZI. Urbain and Deschênes^11^ found that electrical activation of M1 led to a brief inhibition of ZI output neurons for 20-40 ms, which might be expected to facilitate trans-thalamic excitation, but would not explain subsequent suppression of sensory responses. Interestingly, Urbain and Deschênes^11^ were unable to observe disinhibitory effects of motor cortex stimulation on PoM responses, which they attributed to the recruitment of cortico-thalamic fibres from layer VI, and subsequent inhibition of reticular thalamic origin. Indeed, feedforward and feedback inhibition via reticular thalamus would be a parsimonious explanation for the tactile suppression we observe, with inhibition propagated across PoM and VPM via intrathalamic pathways^33^.

The dynamic gating of sensory responses via M1 cortico-thalamic projections bears a resemblance to that shown for S1 layer VI cortico-thalamic projections to VPM^34^. A similar temporal profile of inhibition can be induced by paired whisker stimulation, which may have a strong cortical components^35^, but is also likely to be implemented to some extent within thalamus^36^. This could suggest that some of the circuitry utilised to establish the delayed inhibition could be shared between intramodal and cross-modal computations.

While the experiments exploring the effects of activating M1 NTSR1+ layer VI cortico-thalamic neurons provided clearer evidence for both a trans-thalamic M1-S1 pathway and a thalamic locus of M1-induced tactile suppression, we observed qualitatively similar results with activation of M1 RBP4+ layer V neurons. These layer V and VI neurons have different cortico-cortical and corticofugal projections, which must influence their functional output, and our experimental paradigm is unlikely to be sufficiently refined to resolve the different and/or complementary roles played by subsets of M1 infragranular neurons in somatosensory processing. However, pyramidal tract-projecting neurons in layer V and cortico-thalamic neurons in layer VI are functionally disconnected^13^, and these layers receive different patterns of projections from higher-order cortical structures^37^. Therefore, it is possible that M1 projections could have qualitatively similar effects on sensory processing while being used flexibly during behaviour, with, for example, layer VI coritco-thalamic projections enabling the modulation of somatosensory thalamocortical processing during movement preparation^38^.

The role of the thalamus, and its interaction with cortex, is currently being revisited^17,34,39-42^. By investigating the role of motor corticofugal layers on somatosensory thalamo-cortical processing, we provide evidence of a novel circuitry and role for the thalamus to play in establishing a link between action and sensation.

## Methods

All experiments were approved by the local ethical review committee at the University of Oxford and licensed by the UK Home Office.

### Animals and viral vectors

All experiments were carried out on adult mice bred on a C57BL6/J background. For optogenetic experiments, viral delivery of AAV5-DIO-ChR2(h134R)-eYFP, AAV5-CaMKII-ChR2(h134R)-eYFP constructs were pressure injected into C57BL6J (ChR2), RBP4-cre (DIO-ChR2), or NTSR1-cre (DIO-ChR2) mice.

### Viral delivery and surgery

Mice were intracortically injected with floxed (RBP4-cre and NTSR1-cre mice) or non-floxed (C57BL6/J) AAV5-ChR2-eYFP.

Mice had anaesthesia induced in an induction chamber filled with 4 % isoflurane. Mice were subsequently placed in a stereotaxic frame with a constant flow of 2-4 % isoflurane (adjusted in order to keep breathing rate about 1 Hz) in a 1.5 litres/minute oxygen flow. The isoflurane was scavenged with an active gas scavenging unit. Before any surgical procedures, systemic pain relief was subcutaneously administered with 20 μl buprenorphine (10 μg/ml) and 80 μl Meloxicam (250 μg/ml). For local pain relief 40 μl (2.5 mg/ml) bupivacaine was administered under the scalp prior to incision. A small craniotomy above the two injections was performed. The exposed cranium and dura-mater was kept moist with saline during the entire procedure.

Injections were targeted at left vM1. Two injections of the viral construct were performed. Estimated from bregma, injections were placed at 1.20mm lateral, and 1.00mm and 1.40mm rostral, respectively. Injections were aimed at layer V (RBP4 mice) or Layer VI (NTSR1 mice) and were placed approximately 650-800 microns, and 750-900 microns deep, respectively. 400 nl of the AAV5-ChR2-YFP was delivered manually over 10 minutes at each injection site using a Hamilton syringe. The syringe was left for 2 minutes after injection was finished to avoid the virus flowing away from the injection site. The syringe was then taken slowly out of cortex over 1-2 minutes. Histology confirmed that eYFP-ChR2 was expressed selectively in layer V for RBP4 mice and in layer VI for NTSR1 mice (see Supplementary Figure 2). For post-surgical pain relief, animals were given 80 μl meloxicam the subsequent day. Mice survived and recovered 3-5 weeks for expression of ChR2 to occur.

The electrophysiological experiments were carried out 3-5 weeks after viral injection. All recordings were performed under anaesthesia. Mice were injected with 10ul/g of a mix of medetomidine (100 μg/ml) and ketamine (10 mg/ml) in sterile water (i.e. Ket-Med-Mix). Mice were given additional Ket-Med-Mix if the initial dose did not sufficiently remove the pedal withdrawal reflex. Furthermore, 40 μl bupivacaince were administered under the scalp prior to incision. Top-ups of 50 μl ketamine (10 mg/ml) were given via an intraperitoneal line every 30-60 minutes, or when pedal reflex was strong between recordings. Three hours into the procedure, mice were given half the initial dose of the Ket-Med-Mix in 50 μl top-ups. Mice were kept on a heating blanket during the whole procedure, and body temperature was kept between 38-39 ^o^C. Furthermore, mice were subjected to a constant oxygen flow of 0.5 litre/minute. If the anaesthesia became too light, the oxygen was increased to 1.5 litre/minute and mixed with 1-2 % of isoflurane. Given that isoflurane strongly suppresses cortical activity recordings were paused for 20-30 minutes after the removal of isoflurane, to regain cortical responses. Isoflurane was very rarely used during experiments. Two craniotomies were performed. One exposed the areas of the initial injection site (vM1), and the other exposed areas above vS1 and several thalamic nuclei. At the end of the experiment animals were lethally overdosed with pentobarbital.

### *In vivo* data collection and analysis

#### *In vivo* multielectrode extracellular recording and whole-cell current-clamp recording, and analysis

Whole-cell current-clamp recordings were performed with glass pipettes (3-8 MΩ), pulled from standard borosilicate glass, and filled with a pipette solution containing (in mM): K-gluconate (110), HEPES (40), ATP-Mg (2), GTP (0.3), NaCl (4) and 4 % biocytin (wt/vol) (pH 7.2-7.3; osmolarity 280-290 mosmol/l). The craniotomies were bathed with cortex buffer solution containing (in mM): NaCl (150), KCl (2.5), HEPES (10) (pH 7.4). Recordings were acquired in pClamp 10.0 using a Multiclamp 700B amplifier and DigiData 1440A A/D board (Molecular Devices). Cells were included in the analysis if the action potential amplitude exceeded 50 mV and input resistance was lower than 150 MΩ. The amplitude of whisker-evoked EPSPs were measured relative the membrane potential in the 5 ms prior to stimulation. For responses that evoked action potentials, the EPSP amplitude was estimated from the periods of the response not contaminated by action potentials – action potential onset was defined as the point at which the differentiated membrane potential exceeded the baseline value by 5 standard deviations, and action potential offset as the point at which the differentiated membrane potential returned to zero.

We recorded extracellular activity using linear NeuroNexus probes in a 1×16 configuration with either 50 or 100 micron spacing between electrodes. Extracellular signals were amplified on a TDT RA16PA Medusa preamp and send to a TDT RZ2 BioAmp Processor. Data were acquired using Brainware, and multi-units were detected using a 3 std threshold of the high pass filtered extracellular signal. We recorded from either the somatosensory brain stem (trigeminal nuclei), thalamus (VPM or PoM), or S1 in succession.

Whisker responses were calculated as the average number of spikes in a 45 ms response window (5-50 ms following whisker stimulation onset). This response window captures the peak response of all recorded MUAs across the somatosensory pathway. Due to the variable delay of M1 stimulation induced responses, M1 stimulation induced responses were calculated as the average number of spikes 20 ms around the peak of the M1 stimulation induced response for individual MUAs. Response delays were measured as the time point (1 ms bins) following stimulation (within first 100 ms) where the maximum firing rate was measured.

#### Stimuli

Whisker deflections consisted of a single valley-to-valley cosine wave lasting either 10 or 20 ms. Whisker deflection stimuli were programmed in BrainWare and were delivered by inserting all principal whiskers into a tube attached to a piezoelectric bimorph controlled by a RP2 real-time processor.

For *in vivo* patching (and a subset of extracellular recordings for piloting studies), we performed intracortical microstimulation of M1 using glass pipettes (0.5-1 MΩ) filled with cortex buffer. We stimulated the part of M1 (layer V) that we found to produce retraction of the whiskers when electrically stimulated with train stimulation of electrical pulses.

For all extracellular recordings reported in the main text we performed optogenetic manipulation of M1 activity. This was done by placing a 200 μm fiber-optic above M1 and delivering single pulses 5-10 ms of 1.5-3.6 mW 465 nm light. We targeted the LED in the approximate position we had previously found to elicit retraction of the whiskers during electrical stimulation in other animals.

#### Whole-cell patch-clamp recordings in brain slices

Mice were decapitated under isoflurane anesthesia, and the brains removed in a warm (~32°C) sucrose-based cutting solution, containing (in mM): sucrose (130), NaH_2_PO_4_ (1.25), KCl (3), MgSO_4_ (5), CaCl_2_ (1), NaHCO_3_ (25), D-glucose (15), with pH 7.2-7.4 with when bubbled with carbogen gas (95% O_2_ and 5% CO_2_). Coronal/horizontal slices (350 μm) were prepared using a Vibratome VT1200S (Leica, Germany), transferred to an interface recovery chamber filled with artificial cerebrospinal fluid (aCSF) containing (in mM): NaCl (126), KCl (3.5), NaH_2_PO_4_ (1.25), MgSO_4_ (1), 1 CaCl_2_ (1.2), NaHCO_3_ (26), and 10 glucose, with pH 7.2-7.4 when bubbled with carbogen gas. The slices were maintained at 32-34 °C for 1 hour, before being allowed to cool to room temperature. For recordings, slices were transferred to a submerged chamber, and superfused with carbogenated aCSF heated to 32-34 °C at 2-4 ml/min. Neurons were visualised under infrared oblique illumination (Olympus, BX51WI, 40x water-immersion objective). Whole-cell current-clamp recordings were performed with glass pipettes (5-8 MΩ), pulled from standard borosilicate glass, and filled with a pipette solution containing (in mM): K-gluconate (110), HEPES (40), ATP-Mg (2), GTP (0.3), NaCl (4) and 4% biocytin (wt/vol) (pH 7.2-7.3; osmolarity: 280-290 mosmol/l). Recordings were acquired using a Multiclamp 700B amplifier (Molecular Devices), and digitised using an ITC-18 A/D board (Instrutech). Blue light was delivered via a galvanometer-based movable spot illumination system coupled to the epifluorescence port of the microscope using a single mode fibre (473 nm, 5-25 ms, UGA-40, Rapp OptoElectronic). Stimulation and recordings were controlled via custom-written procedures in Igor Pro (Wavemetrics).

## Data availability

Supporting data are available on request.

## Acknowledgements

This research was supported by a Marie Curie Action Career Integration Grant (303845) and John Fell Fund award to E.O.M., and a Wellcome Trust PhD scholarship to M.L. (105241/Z/14/Z).

## Author contributions

Conceptualization by E.O.M. and A.L.U.; Data collection by M.L., M.C., E.S., A.L., M.C.K., L.B., J.B., and J.P.; Data analysis: M.L.; Writing M.L. and E.O.M; Visualisation by M.L.; Supervision by E.O.M. and A.L.U.

The authors declare no competing financial interests.

